# DistortCorrect Stitcher: Microscopy Image Stitching with Integrated Optical Distortion Correction

**DOI:** 10.1101/2024.10.28.620720

**Authors:** Hu Cang, Huaxun Fan, Sha Sun, Kai Zhang

## Abstract

The rapidly advancing field of imaging-based spatial transcriptomics demands microscopy images with larger fields of view (FOV) and high spatial resolution to accurately and efficiently quantify RNA molecules at the single-cell level. This challenge requires advanced image processing algorithms to seamlessly stitch multiple microscopic images while overcoming the inherent FOV limitations of microscopes. However, image stitching accuracy is often compromised by optical distortions, which are typically uncorrected or require cumbersome hardware-based calibration. Here, we present DistortCorrect Stitcher, a powerful software package that simultaneously estimates and corrects optical distortion during the stitching process. By iteratively integrating image stitching with distortion correction, our method significantly enhances alignment accuracy without relying on external calibration, providing a robust solution for processing spatial transcriptomics data. DistortCorrect Stitcher maintains high stitching fidelity across large tissue areas, ensuring that the molecular data remain reliable and biologically meaningful. This integrated approach has broad implications for improving spatial biology analysis and enabling more accurate high-throughput assays, thereby accelerating advancements in the field.

## 1. Introduction

Microscopy image stitching^1-11^ is essential for extending the field of view (FOV) in high-resolution imaging. Traditionally, expanding the FOV was achieved by using low-resolution optics, which resulted in a loss of spatial detail. However, the advent of spatial transcriptomics demands both high spatial resolution and a large FOV. High-resolution imaging is crucial for resolving individual RNA molecules, while a large FOV is needed to capture the context of entire tissue structures. To meet these demands, multiple high-resolution image tiles are captured and subsequently stitched together.

Microscopy stitching differs fundamentally from image stitching in other fields, such as photography, where the primary goal is visual consistency. In microscopy, quantitative accuracy of biological features is paramount—misalignments or distortions between tiles can lead to significant errors, impacting downstream data interpretation. Spatial transcriptomics^12-16^, in particular, requires extremely precise stitching, as these methods rely on detecting individual RNA molecules, visible as diffraction-limited puncta. Misalignments between image tiles can cause RNA signals to be misplaced, leading to errors in quantification and erroneous data interpretation. Unfortunately, mechanical errors in microscope stage movements often introduce inaccuracies in the positioning of each tile used for stitching. Therefore, routine microscopy procedures, which involve moving the stage holding the sample to preset coordinates and using those coordinates to stitch the final images, are prone to positioning errors that affect image alignment, necessitating more advanced correction techniques.

To address these challenges, computational tools such as M2Stitch^8^, AstroPath^9^, MIST^11^, ASHLAR^7^, and BigStitcher^10^ have been developed to realign neighboring tiles and stitch them together. These tools use overlap regions between neighboring tiles to calculate their relative positions and assemble those tiles into a complete image. Such stitching tools are indispensable for advancing spatial transcriptomics, ensuring both spatial resolution and coverage.

However, optical distortions^17^ pose a significant challenge to achieving precise image stitching. These distortions are caused by intrinsic aberrations in microscope lenses, resulting in nonlinear warping of images, particularly near the edges. Figure 1A illustrates barrel and pincushion distortions, the two types of optical distortions. Imagine a camera attempting to image a flat grid, represented by blue lines. In an ideal case, without distortion, the captured grid lines would remain flat. However, the magnification across the FOV is not uniform due to lens imperfections, leading to differential magnifications along the radius of the FOV, which in turn distorts the grid lines in the image (black lines).

**Fig. 1.**
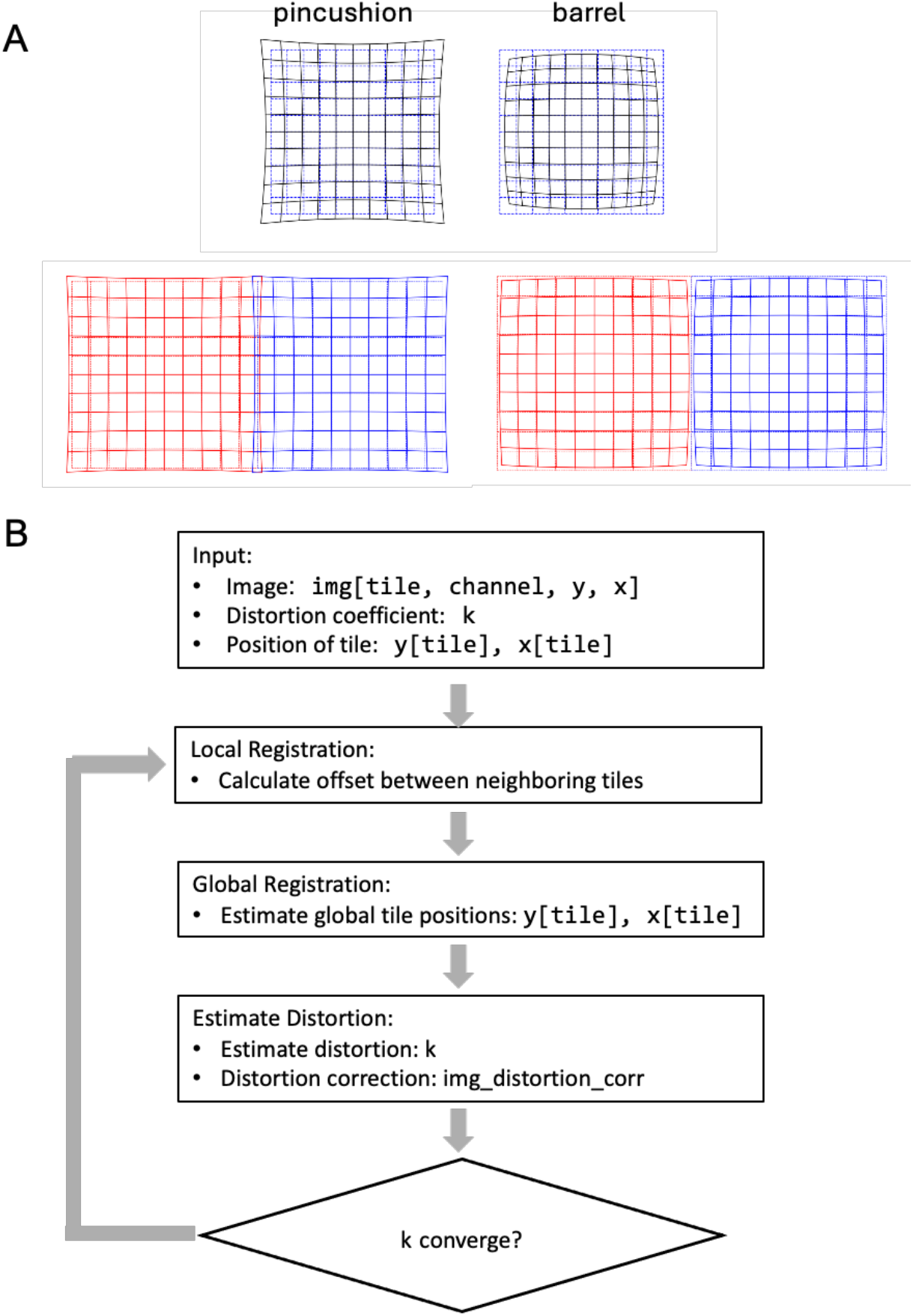
Image Stitching and Distortion Correction Workflow. **(A)** Two examples of optical distortions: **pincushion distortion** (left) and **barrel distortion** (right). These distortions show how images get warped due to optical imperfections, with pincushion pulling the grid towards the center and barrel pushing it outward. Below are two grids representing the tiled images, with the left tile (red) and right tile (blue) used in the registration process. **(B)** Flowchart illustrating the iterative process of image stitching with distortion correction. The input includes the image tiles (img[tile, channel, y, x]), distortion coefficient (k), and the tile positions (y[tile], x[tile]). The process begins with local registration, where offsets between neighboring tiles are calculated. In global registration, the global positions of the tiles are estimated and updated. The distortion estimation step uses overlapping regions between tiles to estimate the distortion coefficient (k) and generate the distortion-corrected image (img_distortion_corr). The process iterates, refining the registration and distortion correction until k converges.

In spatial transcriptomics, the imaged coordinates of an RNA molecule, denoted as *x’*, are related to its true coordinates *x* by a radial distortion polynomial of the form *x*′ *= x*(*k*_1_*r*^2^ + *k*_2_*r*^4^ + *k*_3_*r*^6^), where *r* is the distance from the optical center, which often coincide with the center of an image tile, and *k*_*1*,*2*,*3*_ are distortion coefficients. Even in high-quality microscope objectives lens, distortions ranging from 0.1% to 0.5% are common. Given that a typical image may span an area of 500μm, these distortions can cause shifts near the edges that exceed the size of an individual RNA spot ∼0.5μm, making accurate stitching highly challenging.

The nature of optical distortions is dictated by the optical characteristics of the lens. As shown in Figure 1A, two neighboring tiles, represented by red and blue grids, exhibit the same type of distortion (either barrel or pincushion). As a result, the overlapping areas of these neighboring tiles warp in opposite directions—toward the center of each tile. Conventional stitching algorithms, which rely solely on translation to align tiles, cannot correct for these opposing warps. Consequently, RNA molecules in the overlapping regions appear as misaligned spots between the neighboring tiles, preventing seamless alignment across all overlap areas. Without accurate distortion correction, this misalignment can lead to inaccurate RNA molecule quantification and difficulties in downstream analysis. One possible approach to mitigate distortion is to reduce the FOV, as distortions tend to be more severe near the edges. However, this requires taking more images to cover the same area of a sample and significantly sacrifices throughput, which may not be feasible for high-throughput spatial transcriptomics. Existing image stitching algorithms, such as M2Stitcher, MIST, and BigStitcher, can align neighboring tiles but provide limited capabilities for correcting optical distortions. AstroPath offers distortion correction, but it is performed separately from the image stitching process. Without hardware-based calibration, estimating distortions requires comparing overlapping regions between neighboring tiles. At the same time, these overlap regions are computationally determined based on image similarity, making distortion correction and stitching highly interconnected tasks.

Here, we present DistortCorrect Stitcher, an effective software package that simultaneously corrects optical distortions and stitches image tiles. By iteratively integrating image stitching with distortion correction, our method significantly enhances alignment accuracy without relying on external calibration, providing a robust solution for processing spatial transcriptomics data. This integrated approach maintains spatial accuracy across large tissue areas, ensuring that the molecular data remain robust and biologically meaningful. This tool also has broad implications for improving spatial biology analysis by enabling more accurate high-throughput assays, thereby accelerating advancements in the field.

## 2. Method

### 2.1 Workflow Overview

DistortCorrect Stitcher introduces a novel image stitching method that integrates distortion correction directly into the stitching process, significantly enhancing the accuracy and quality of microscopy image mosaics. Traditional approaches typically handle image stitching and distortion correction separately, where distortion calibration is performed before aligning the image tiles, often requiring additional hardware-based calibration tools. In contrast, our method integrates distortion correction into the stitching workflow, effectively estimating and correcting optical distortion during the iterative alignment of image tiles.

The workflow of DistortCorrect Stitcher is illustrated in Figure 1B and consists of the following steps: (1) acquisition of multiple overlapping image tiles using high-resolution microscopy; (2) local alignment of overlapping regions; (3) global optimization to ensure consistency across all tiles; (4) estimation and correction of optical distortion; and (5) iterative refinement of optical distortions and global coordinates until convergence. By integrating alignment and distortion correction processes iteratively, DistortCorrect Stitcher achieves superior stitching accuracy, effectively overcoming challenges posed by optical aberrations.

Each image tile, *img*[*tile*], is captured with approximately 10% overlap with neighboring tiles. It is worth noting that in other stitching algorithms, the overlap is often required to between 15–20%, which reduces experimental throughput. Our approach reduces the required overlap while maintaining high-quality stitching. This reduced overlap contributes to improved efficiency, making the stitching process more suitable for high-throughput spatial transcriptomics experiments. By iterating the stitching and correction steps, the method converges to an optimal solution that corrects both local misalignments and global distortions, ensuring robust and biologically accurate data.

### 2.2 Local Alignment Methods

Microscopes record the coordinates of each image tile, but these recorded coordinates are often not sufficiently accurate for precise alignment. Therefore, we initially use the recorded coordinates as a rough guide to estimate the overlap regions between neighboring tiles. The relative positions between neighboring tiles are subsequently refined based on these estimated overlap regions.

Consider two neighboring tiles, *img*[*tile*_*i*_] and *img*[*tile*_*j*_]. To align these tiles, we first identify the overlap regions, defining a local region in each tiles as *overlap_img*[*tile*_*i*_] and *overlap_img*[*tile*_*j*_]. We estimate relative local position between the two tiles as *P*_*ij*_, by maximizing the similarly of the overlapped regions from the two tiles. *P*_*ij*_ is pairwise, for tiles with no overlapping, *P*_*ij*_ *=* 0.

In spatial biology, phase correlation^18^ is commonly used for image registration to calculate *P*_*ij*_. Phase correlation computes the cross-correlation between pixel intensities in the overlapping regions of neighboring tiles, which can be represented as *Corr*(*overlap_img*[*tile*_*i*_], *overlap_img*[*tile*_*j*_]). The algorithm shifts one tile relative to the other to maximize this cross-correlation value, which should reach a maximum when the tiles are correctly aligned. However, phase correlation is prone to instability, often producing multiple correlation peaks, which makes determining the correct alignment challenging.

To overcome these limitations, we employ mutual information (MI)^19^ as an alternative approach for image registration. Mutual information^20-22^ measures the amount of information shared between the overlapping regions of two images, which makes it robust to intensity variations and less sensitive to noise. MI is widely used in the medical imaging community for aligning images acquired from different modalities, as it effectively handles complex relationships between image intensities.

Mutual information (MI) between two regions *A* and *B* images quantifies the amount of information one can derive about one region based on the other, assessing the statistical interdependence between the intensities of the two. Normalized mutual information is calculated using the following formula: *MI*(*i, j*) *=* 2(*H*(*i*) + *H*(*j*) ™ *H*(*i, j*))/(*H*(*i*) + *H*(*j*)). Here, *H*(*i*) is the entropy of *overlap_img*[*tile*_*i*_], *H*(*j*) is the entropy of *overlap_img*[*tile*_*j*_], and *H*(*i, j*) is the joint entropy of *overlap_img*[*tile*_*i*_] and *overlap_img*[*tile*_*j*_]. The entropy H of an image could be calculated from the pixel intensity. The normalized mutual information is in the range of 0 (no correlation) and 1 (exactly the same). When mutual information is high, it implies a strong alignment between the pixel intensities of the two tiles.

In our approach, we adapt a Fast Fourier Transform (FFT) based algorithm^19^ to calculate MI and estimate the relative positions *P*_*ij*_ for all neighboring tile pairs. By maximizing MI, we determine the optimal alignment that ensures consistent overlap between neighboring tiles, even in the presence of intensity variations or noise. This makes MI a reliable method for spatial transcriptomics applications, where image quality may vary across different tiles due to biological or experimental factors.

### 2.3 Global Alignment Optimization

Each image tile may have multiple neighbors, and these neighboring tiles also overlap with each other. Therefore, after determining the local coordinates for each pair of neighboring tiles *P*_ij_, we must compute a consistent set of global coordinates for each tiles. To achieve this, we employ a global optimization approach that minimizes discrepancies across all local alignments. Specifically, we minimize the following objective function: ∑_*ij*_ (*P*′_*ij*_[*x, y*] ™ *P*_*ij*_)^2^ to find the global coordinates of all image tiles [*x, y*]. Here, *P*′_*ij*_ is the relative position calculated from the global coordinates of tile *i* and *j*, and *P*_*ij*_ is calculated from local alignment via maximizing mutual information between the overlapping regions of tile *i* and *j*. By minimizing this objective function, we ensure that the global coordinates are consistent with the estimated local alignments, resulting in an accurate reconstruction of the entire field of view.

We solve this optimization problem using an iterative algorithm that refines the global positions until discrepancies between local and global alignments are minimized. This iterative approach ensures accurate and consistent alignment of all image tiles, producing high-quality stitching results without the need for external calibration. The global optimization step is essential for maintaining spatial consistency across the entire stitched image, particularly in scenarios where local alignment alone may introduce cumulative errors.

### 2.4 Distortion Correction

After determining the global coordinates of each image tile, we correct optical distortions from the overlapping regions between all neighboring tiles. Optical distortions inherent in microscope optics are modeled using a polynomial function of the radial distance from the optical axis. This model captures first-order, second-order, and third-order distortions through coefficients *k*_1_, *k*_2_, *k*_3_. The relationship between the distorted and true coordinates is given by: *x* ′ *= x*(*k*_1_*r*^2^ + *k*_2_*r*^4^ + *k*_3_*r*^6^), where **r** is the distance from the optical center.

To evaluate the similarity between neighboring tiles, we calculate the normalized correlation (ncc)^23^ of each overlapping region. If two neighboring tiles are perfectly corrected for distortion and properly aligned, their overlapping pixels should yield a normalized correlation value of 1. Since optical distortion is an intrinsic property of the microscope optics and consistent across all image tiles, we fit a global set of distortion coefficients (*k*_1_, *k*_2_, *k*_3_) by minimizing the sum of normalized correlation across all overlapping regions from all neighboring tiles. The total normalized correlation is expressed as: ∑ *ncc*(*i, j*), where *ncc*(*i, j*) represents the normalized correlation between the overlapping regions of neighboring tiles *i* and *j*. The global optimization ensure that all overlapping regions are optimally corrected for distortion, providing consistency and improved alignment accuracy across the entire image set.

### 2.5 Iterative Refinement

After correcting for distortion, we used the corrected images to recalculate both their local and global coordinates. This iterative process continued until we obtained a converged estimation of the distortion coefficients. This correction is performed in conjunction with the stitching process, rather than separately, ensuring that the distortion model evolves as the tiles are aligned. With each iteration, both the alignment and distortion corrections improve, converging towards an optimal solution where the entire mosaic is accurately stitched with minimized distortion effects.

## 3. Result

### 3.1 Distortion Correction and Image Stitching of Two Imaging Tiles

We first demonstrate the effectiveness of DistortCorrect Stitcher on two neighboring imaging tiles. Figure 2 shows an image of mouse liver tissue stained for RNA Malat1 using fluorescence in situ hybridization (FISH). The image was captured using a Leica microscope equipped with a 20X objective lens with a numerical aperture (NA) of 0.8. The FISH probe uses Alexa488 fluorescence dye, and the puncta in the images represent Malat1 RNA molecules. The two imaging tiles have a 10% overlap region. Each tile measures 2048 × 2048 pixels, with each pixel corresponding to 0.33 micrometers. The approximate spot size of each punctum is 1.22 × 488/0.8 = 0.7*μm*, implying that the accuracy of image registration must be below 0.7 micrometers to avoid counting the same RNA molecule twice.

**Fig. 2.**
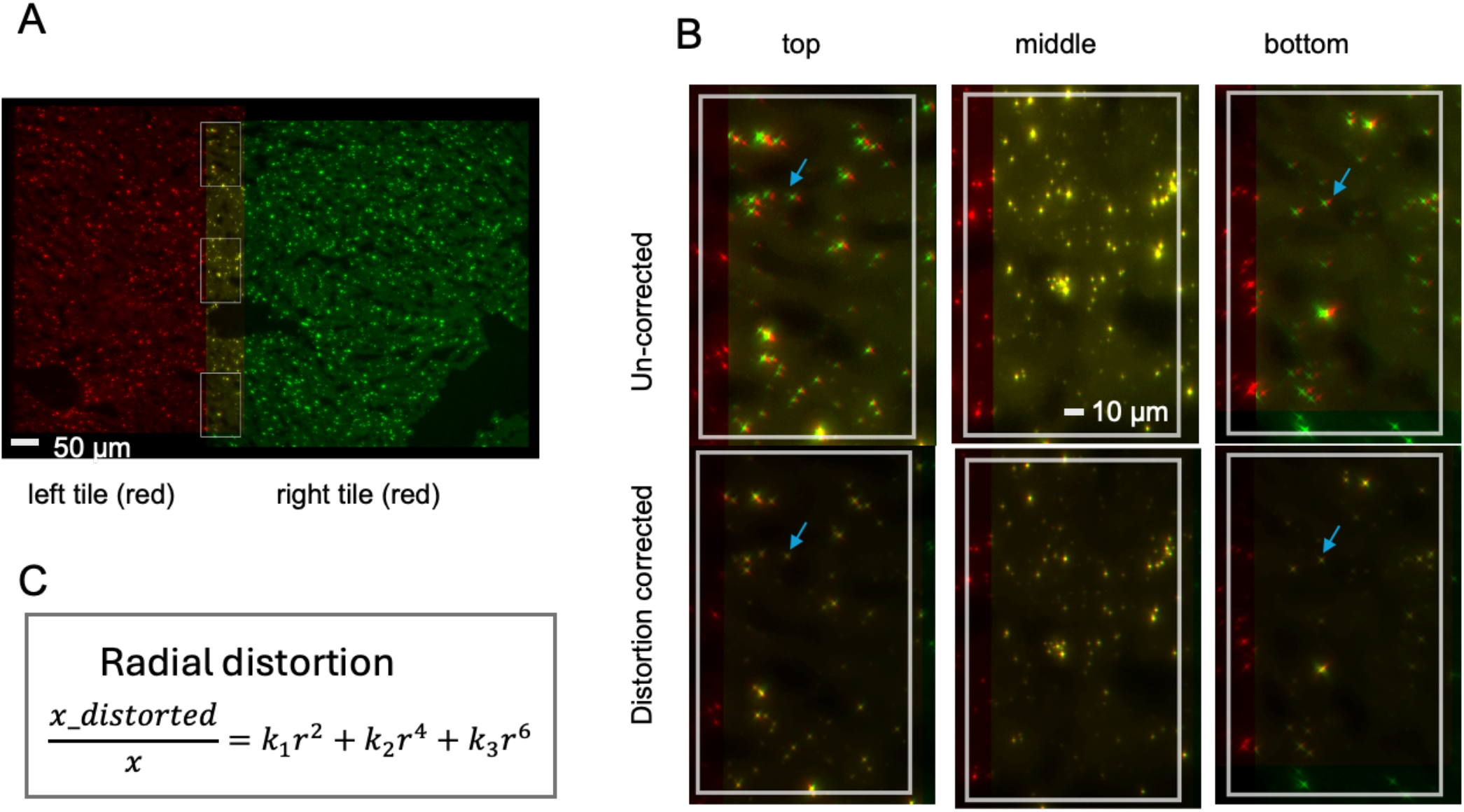
Misalignment of Image Tiles in Stitching and Improved Alignment after Correction. **(A)** Overview of image tiling in mouse liver tissue, where each punctum represents RNA fluorescence in situ hybridization signals. The red and green channels represent the left and right image tiles, respectively. Overlapping RNA FISH signals from the left and right tiles appear yellow. The white box marks a zoomed-in region spanning two neighboring tiles. Scale bar = 50 µm. **(B)** Zoomed-in view of the stitched image without distortion correction, demonstrating clear misalignment in the top and bottom regions (blue arrows), particularly between puncta representing FISH signals of RNA molecules. Misalignment is less pronounced in the center. **(C)** Zoomed-in view after distortion correction, showing improved alignment across the entire region. The radial distortion is characterized by a polynomial, and fitting to the first-order term, *k*_1_, is sufficient to achieve good distortion correction. The scale bar in the zoomed-in views represents 20 µm.

To better visualize the stitching, we show the two tiles in different colors—red and green. FISH signals in the overlap region appear in both the red and green channels. If the stitching is successful, the red and green signals of the same molecule will overlap perfectly, producing yellow puncta. If not properly stitched, the same RNA molecule will appear as two distinct puncta in red and green, respectively. Three zoomed-in regions are shown in Figure 2B.

The top panel of Figure 2B shows the result of stitching without distortion correction. The effect of optical distortion is clearly visible. In both the top and bottom zoomed-in regions, the alignment is poor, and each RNA punctum appears as two distinct dots in red and green, while the middle region is aligned well, forming yellow puncta. Importantly, in both the top and bottom regions, the green puncta consistently appear to the left of the red puncta, whereas in the middle region they overlap perfectly, forming yellow puncta. This demonstrates that the images are warped: the top and bottom areas, which are further from the center of each tile, exhibit greater warping compared to the middle region. This is characteristic of optical distortion. The misalignment due to optical distortion presents significant challenges for accurate stitching. Since spatial transcriptomics relies on these puncta to count RNA molecules in each cell, distortion in regions with pronounced misalignment can result in inaccurate RNA quantification.

The lower panel of Figure 2B shows the same zoomed-in regions after using DistortCorrect Stitcher, which simultaneously corrects optical distortion and stitches the image tiles together. It is clear that all three regions are aligned well, with each RNA molecule appearing as a single yellow punctum, indicating perfect alignment. The blue arrows highlight the same RNA molecules in both the distortion-uncorrected (top panel) and corrected (bottom panel) images, showing that distortion correction results in improved alignment.

Figure 2C illustrates the formula used for image distortion correction. The distortion correction employs only the first-order distortion coefficient *k*_1_, which is sufficient to achieve acceptable correction for image stitching. Therefore, we recommend starting with *k*_1_ only. Our software also supports higher-order distortion corrections using *k*_2_ and *k*_3_, but starting with *k*_1_ is advisable.

This result with two tiles demonstrates the challenge of achieving high spatial resolution to accurately count sub-micrometer-sized RNA molecules while maintaining a large field of view to profile larger tissue areas. High-accuracy image processing algorithms are essential for stitching these images, as they must overcome the inherent optical aberrations of the microscope. Without addressing optical distortion, artifacts may arise, leading to inaccuracies in RNA quantification. DistortCorrect Stitcher, which estimates and corrects optical distortions concurrently with image stitching, effectively overcomes these challenges. It preserves alignment fidelity near the boundaries of image tiles, thereby achieving both high resolution and high throughput for spatial transcriptomics. Importantly, DistortCorrect Stitcher does not require any prior calibration of optical distortion—it estimates the distortion directly from the images, making it convenient for practical applications.

### 3.2 Distortion Correction and Image Stitching of 28 Imaging Tiles

We next demonstrate the effectiveness of distortion correction and image stitching for an entire tissue sample consisting of 28 imaging tiles. This section showcases the efficiency of the global registration capability of DistortCorrect Stitcher. Unlike the previous example with only two imaging tiles, this scenario requires global coordinates for each tile, as most imaging tiles have at least four neighboring tiles, resulting in multiple pairs of local alignments. However, each imaging tile has only one global coordinate, leading to an overdetermined problem.

AstroPath^9^ uses a spring-like model to derive global coordinates from local registrations, while M2Stitch^8^ and MIST^11^ utilize a graph-based minimal spanning tree algorithm. We adopted an approach similar to AstroPath’s. Specifically, 27 global coordinates (for 28 tiles) are treated as fitting parameters, with the position of the first imaging tile arbitrarily chosen as the reference. From these 27 coordinates [*x, y*], we calculate the local relative positions among neighboring tiles, denoted as *P*′_*ij*_[*x, y*]. If two tiles *i* and *j* are not neighbors, then *P*′_*ij*_ [*x, y*]=0. We also have *P*_*ij*_ calculated from local registration. By minimizing the L2-norm ∑_*ij*_ (*P*′_*ij*_ [*x, y*] − *P*_*ij*_)^2^, we determine the best global coordinates [*x, y*].

Once the global registration step is complete, we recalculate the overlapping regions of neighboring tiles and use this information to perform distortion correction. This iterative procedure, illustrated in Figure 1, allows for simultaneous recursive distortion correction and image stitching across all 28 imaging tiles.

Figure 3A provides an overview of the entire stitched image. The white grid lines, highlighted by blue arrows, represent the overlapping regions between neighboring tiles, covering approximately 10% of each tile’s area. Tiles that are not at the boundary generally have at least four neighbors. To evaluate the effectiveness of the image stitching, we selected a region near the boundary of two tiles and zoomed in, as shown in Figures 3B and 3C. Figure 3B displays the stitching result without distortion correction, while Figure 3C shows the result with both distortion correction and image stitching applied. To create the final stitched image, we selected the center of the overlap region as a dividing line: the left half of the region uses the left tile, and the right half uses the right tile. This dividing line is highlighted as a white dotted line in Figures 3B and 3C.

**Fig. 3.**
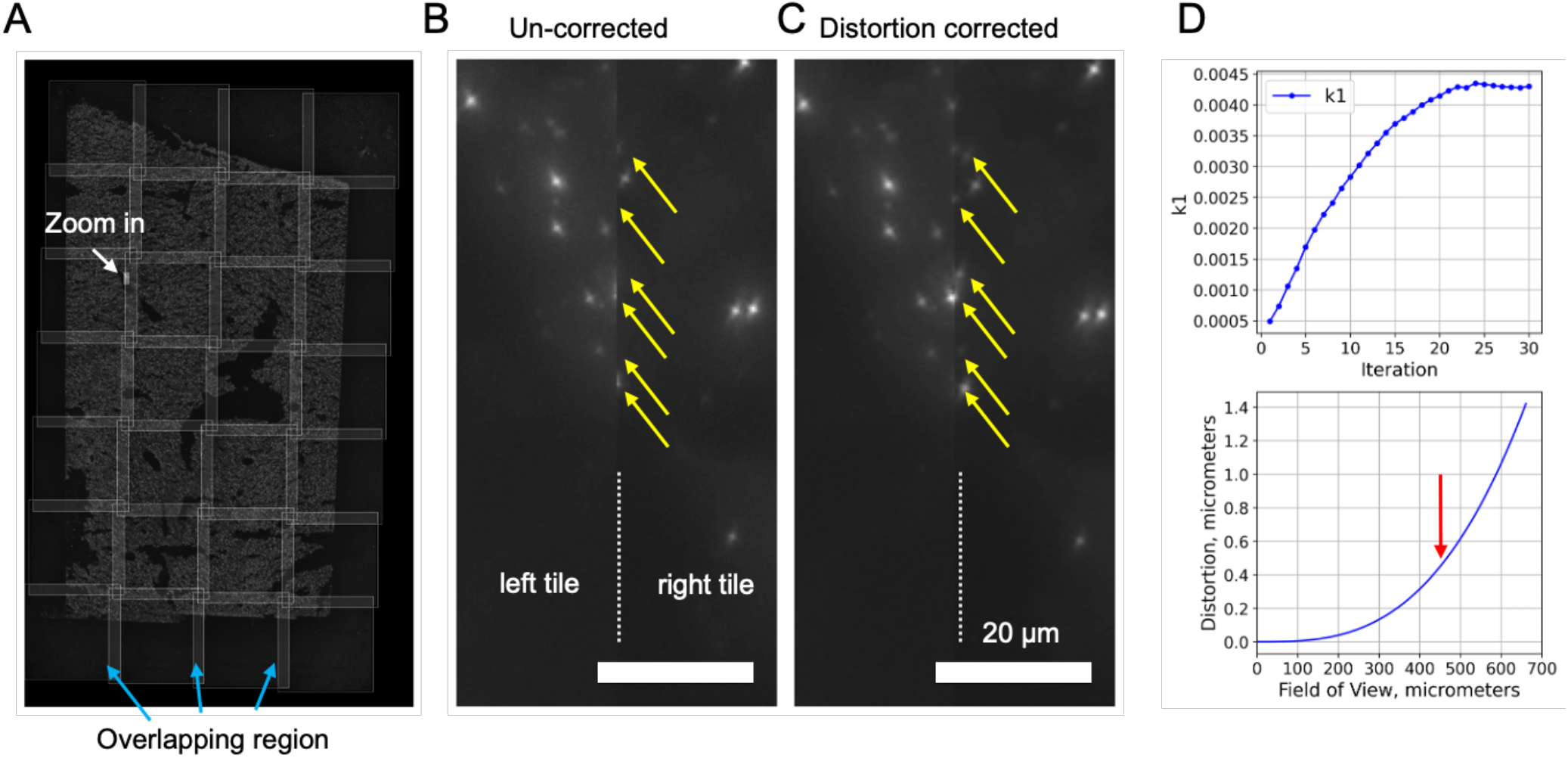
Global Distortion Correction and Image Stitching. **(A)** Overview of the tiled image from mouse liver tissue. The grid represents overlapping areas (∼10%) among adjacent image tiles, with blue arrows indicating regions of overlap. The white box highlights the zoomed-in region displayed in **(B)**. Magnified view of the boundary between the left and right tiles. Yellow arrows indicate the misalignment of fluorescent signals at the boundary when images are registered globally but without distortion correction. The scale bar represents 20 µm. **(C)** Concurrent distortion correction and global image registration correct the misalignments of RNA FISH signals. **(D)** Plot of the distortion correction parameter *k*_1_ over iterative fitting, with the optimal value reached around iteration 25. The bottom panel displays the distortion magnitude as a function of the field of view, illustrating how distortion increases with distance from the center. The red arrow indicates where the distortion reaches 0.5 µm, the size of a diffraction-limited RNA spot.

The difference between the results with and without distortion correction is clearly visible. Without distortion correction, many RNA molecules are missing (highlighted by yellow arrows in Figure 3B), whereas they are well preserved after distortion correction (Figure 3C). These missing molecules would introduce significant artifacts in RNA quantification.

In the upper panel of Figure 3D, we plot the distortion coefficient *k*_1_ against the iteration number. The value of *k*_1_ converges after approximately 22 iterations, suggesting that we could limit the number of iterations to about 22 cycles. Our software also outputs the *k*_1_ parameter, allowing users to monitor the convergence of the iterations.

To further illustrate the impact of distortion on throughput for spatial transcriptomics, we plotted the distortion in micrometers against the size of the field of view (FOV) given the measured *k*_*1*_ parameter in the lower panel of Figure 3D. The distortion, without correction, follows a quadratic form. We set a tolerance threshold of 1 micrometer, beyond which spatial transcriptomics would likely lead to misidentification. Due to the opposite direction of distortion between neighboring tiles—towards the center of each tile—the effective distortion doubles, making the tolerance per tile 0.5 micrometers, as indicated by a red arrow. Based on this, we determine that, with the Leica objective lens used in this experiment, the maximum usable FOV without distortion correction is approximately 450 micrometers.

With distortion correction, the FOV can be extended to the entire 2048 × 2048 pixel area, which corresponds to approximately 660 × 660 micrometers and is limited by the size of the camera. It is possible that DistortCorrect Stitcher could extend the FOV even further. Nevertheless, the maximum FOV with distortion correction is 215% greater compared to that without correction. This improvement effectively doubles the throughput of image acquisition, representing a significant acceleration of the experiment and an enhancement in the accuracy of the results.

### 3.3 Distortion Correction and Image Stitching of 28 Imaging Tiles in Four Colors

Next, we demonstrate the effectiveness of distortion correction in multi-color images. Spatial transcriptomics uses different colors to label different RNA molecules. In this pilot experiment, we used probes targeting different regions of Malat1 RNA with three different colors—Alexa488, Alexa561, and Alexa647. We also stained the nuclei of the cells with DAPI. Thus, the entire image of the mouse liver tissue consists of 28 tiles across four colors.

To stitch these tiles together, we must account for chromatic differences in optical distortions among the different colors. The broad wavelength range from DAPI (about 461 nm) to Alexa647 (approximately 670 nm) spans around 200 nm. To put this in perspective, the difference in wavelengths is roughly half of the wavelength of DAPI. If the optical system were linearly scaled, key parameters such as focal length and magnification would differ by up to 50%. Although modern lenses are designed to minimize these differences through various compensations, chromatic differences are still non-negligible.

To accommodate these chromatic differences, we introduced multi-color channel functionality into DistortCorrect Stitcher. Our software can, in principle, handle an arbitrary number of color channels. Figure 4A shows the workflow, with the differences between the single-color (monochrome) and multi-color versions highlighted in red.

**Fig. 4.**
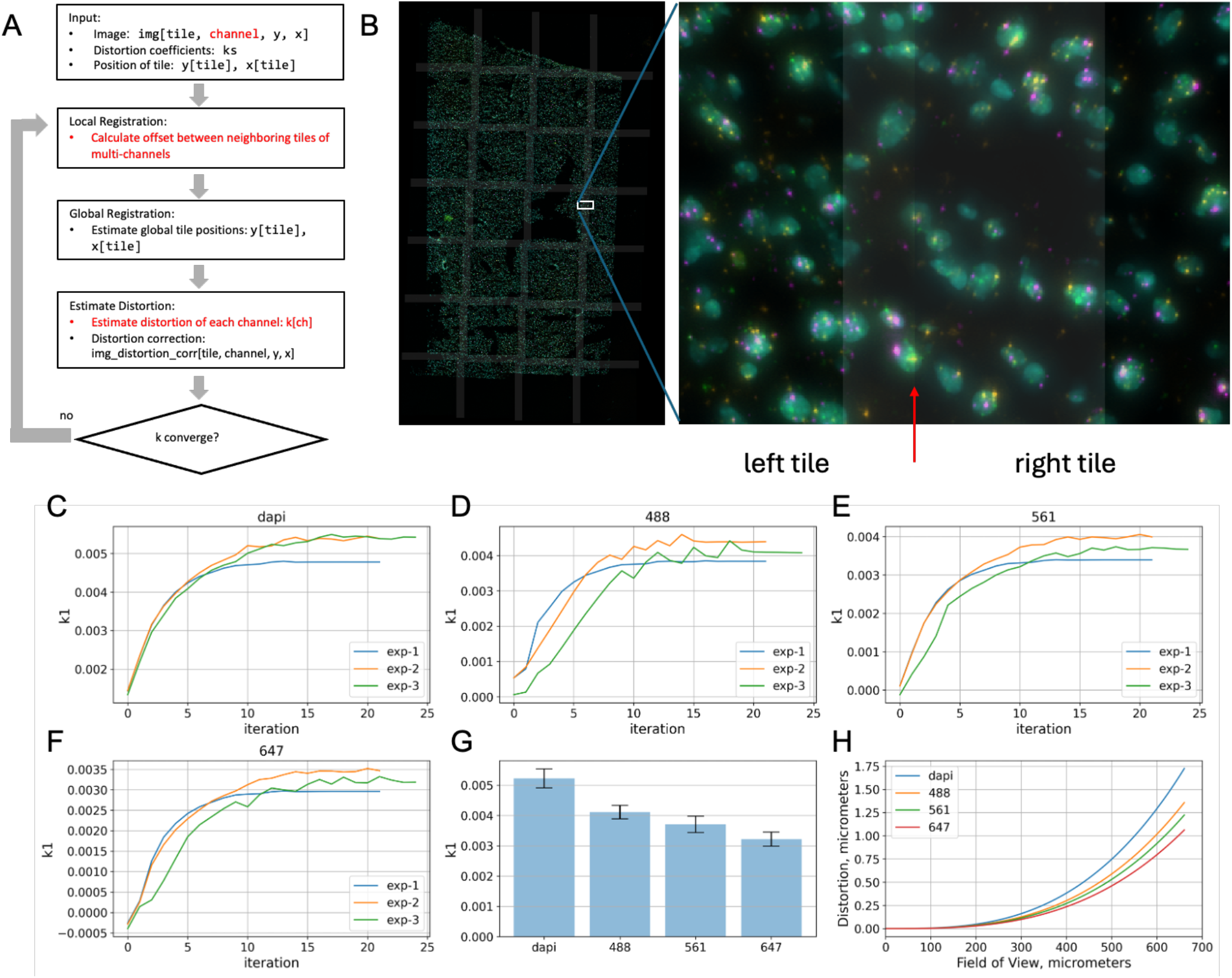
Multi Color Image Stitching and Distortion Correction. **(A)** Flowchart of the DistortCorrect Stitcher algorithm for multi-color imaging. The key steps highlighted in red indicate the modifications for handling multiple channels: calculation of offsets between neighboring tiles using multi-color images and estimation of distortion for each channel *k*_1_. **(B)** Overview of the stitched image of mouse liver tissue, labeled with multiple colors: DAPI (nuclei), Alexa488, Alexa561, and Alexa647 for different regions of Malat1 RNA. The overlapping regions of neighboring tiles are marked with semi-transparent white grids. The are marked with a white box is zoomed-in and shown in the right panel. The red arrow highlights the division between left and right tile. **(C-F)** Plots showing the convergence of the distortion coefficient *k*_1_ for each color channel (DAPI, 488, 561, 647) across three independent experiments (exp-1, exp-2, exp-3). The convergence of *k*_1_demonstrates the iterative correction of distortions over 20-25 cycles. **(G)** The distortion coefficients *k*_1_ for each color channel after convergence. The trend shows higher distortion at shorter wavelengths and lower at longer wavelengths. **(H)** Distortion across the field of view (FOV) in micrometers for all four color channels. The distortion increases with distance from the center of the FOV, and the difference between channels (DAPI, 488, 561, 647) becomes more pronounced near the edges, indicating chromatic differences.

First, when calculating local alignment between neighboring tiles, we modified the mutual information-based local alignment algorithm^19^ to incorporate multiple colors for alignment. Specifically, the mutual information between two images is calculated across all color channels instead of just a single color. Therefore, the quality of local alignment is evaluated comprehensively across all colors. Second, after global registration, distortion correction is performed separately for each color channel. Hence, each color channel has its own set of distortion coefficients *k*_1_. After that, the same iterative procedure is used to perform concurrent image distortion correction and stitching.

Figure 4B shows an overview of the stitched image of mouse liver tissue, labeled with multiple colors: DAPI (nuclei), Alexa488, Alexa561, and Alexa647 for different regions of Malat1 RNA. The overlapping regions of neighboring tiles are marked with semi-transparent white grids. A white box marked an area that is zoomed-in and shown in the right panel. A red arrow highlights the dividing line between left and right tiles in the zoomed in region. The two tiles stitch together seamlessly across all color channels.

Figure 4(C-F) shows the convergence of the distortion coefficients *k*_1_ for each color channel. We observed that convergence is faster—within about 15 cycles—in the multi-color case compared to the single-color case of the same sample. This was independently verified across three different samples, each exhibiting similar convergence behavior.

Figure 4(G) shows the measured distortion coefficients for all four color channels, with error bars representing standard deviations from the three different samples. A significant trend can be observed: distortion is more pronounced at shorter wavelengths and less so at longer wavelengths.

To further illustrate the impact of chromatic differences on distortion, we plotted the distortion in micrometers as a function of the field of view (FOV) for all four colors in Figure 4H. The curves for the four colors diverge near the edge of the FOV, indicating that RNA puncta of different colors could separate and become distinct near the FOV boundary. This observation is particularly important for spatial transcriptomics because current imaging-based spatial transcriptomics relies on multi-color and multi-round imaging to encode different RNA molecules. Each RNA molecule appears in a different color in each imaging round, creating a unique code, such as round 1 (Alexa488), round 2 (Alexa561), round 3 (Alexa488), etc. This temporal coding allows thousands of RNA molecules to be encoded using a limited number of color channels. Our results demonstrate that chromatic differences, if not corrected properly, could cause the same RNA puncta to appear as different signals, leading to incorrect decoding results.

## 4. Discussion

Optical distortion significantly impacts both the accuracy and throughput of image stitching in spatial transcriptomics. Without proper correction, distortion causes misalignment between neighboring image tiles, particularly near the edges of the field of view (FOV). In multi-color imaging, this can lead to the same RNA molecule being misrepresented and decoded differently, compromising data quality and resulting in significant inaccuracies in RNA quantification. Furthermore, optical distortion limits the usable FOV, reducing experimental throughput and efficiency.

Our proposed method, DistortCorrect Stitcher, concurrently estimates optical distortion and performs image stitching, overcoming these challenges by iteratively refining local and global alignments alongside distortion correction. By preserving alignment fidelity across all regions of the imaging field, even near tile boundaries, our method effectively expands the usable FOV— demonstrated by a significant 215% increase in throughput in our experiments.

This increase in throughput, coupled with reduced errors in RNA quantification, directly accelerates the progress of spatial transcriptomics research, facilitating the high-resolution profiling of larger tissue samples with fewer resources. Moreover, our approach eliminates the need for additional hardware-based calibration, making it both accessible and convenient for researchers. The ability to account for chromatic differences in multi-color imaging channels ensures consistent accuracy across all colors, enhancing the reliability of decoding multiplexed RNA signals.

The broader implications of our work are substantial. By providing a robust and efficient tool for both distortion correction and image stitching, DistortCorrect Stitcher paves the way for more accurate, high-throughput spatial transcriptomics, empowering researchers to gain deeper insights into tissue architecture and gene expression at unprecedented scales. Ultimately, this innovation will contribute to advancing our understanding of complex biological systems, aiding in the discovery of novel biomarkers, and opening up new avenues for precision medicine. DistortCorrect Stitcher represents a step forward, enabling the future of spatial biology with unparalleled accuracy and efficiency.

## Experimental Methods

Fresh-frozen mouse liver slices (approximately 10 micrometers thick) were purchased from Zyagen (San Diego) and mounted on 1 mm positively charged glass slides. The samples were fixed in 4% paraformaldehyde (PFA) in phosphate-buffered saline (PBS) overnight. After washing in PBS, DNA nanoballs were generated using DNA probes targeting Malat1 RNA following the RollFISH^16^ protocol. Specifically, a set of 40 DNA probes targeting different regions of Malat1 RNA were hybridized to the sample in 2x saline sodium citrate (SSC) buffer supplemented with 20% formamide. After washing in 2x SSC, 61 padlock probes were hybridized to the initial probes in a similar buffer. The padlock probes were then circularized via ligation using T4 ligase, and Phi29 polymerase was used to elongate the first probes using the circularized padlock probes as a template, forming DNA nanoballs. Three different fluorescence in situ hybridization (FISH) probes labeled with Alexa488, Alexa561, and Alexa647 dyes (ThermoFisher) were then hybridized to the DNA nanoballs. For multi-color experiments, the padlock probes were divided into three groups, each hybridized with a FISH probe of one of the three colors, respectively. Imaging was performed using a Leica fluorescence microscope equipped with a 20x objective lens (0.8 NA). Images were captured using an sCMOS camera (Hamamatsu). To cover the entire tissue, 28 tiles were imaged, with each tile overlapping approximately 10% with neighboring tiles.

## Software Availability

DistortCorrect Stitcher, written in Python, will be published on GitHub after peer review. It is also available upon request.

## Acknowledgements

This work was made possible in part through access to the UCI Optical Biology Core Facility, a shared resource supported by the Cancer Center Support Grant (CA-62203).

